# Bridging The Evolving Semantics: A Data Driven Approach to Knowledge Discovery In Biomedicine

**DOI:** 10.1101/2022.09.05.506661

**Authors:** Kishlay Jha

## Abstract

Recent progress in biological, medical and health-care technologies, and innovations in wearable sensors provide us with unprecedented opportunities to accumulate massive data to understand disease prognosis and develop personalized treatments and interventions. These massive data supplemented with rapid growth in computing infrastructure has enabled bio-medical researchers to perform more comprehensive experiments and detailed case-studies. At the same time, performing these experiments are not only monetarily expensive but also time consuming. Thus, there is a growing need to provide tools to the researchers that will allow them to pose queries that will assist them in focusing on interesting “hypotheses”. However, such a tool would require capabilities to derive inferences based on existing known relationship between medical concepts. In this paper, we tackle this problem as one of non-factoid question answering wherein we try to answer the user-post questions by leveraging both authoritative as well as social media posts. While the former provides us with well knowledge on well researched topics, the latter provides us with real-time feedback on variety of topics like adverse drug effect (ADE), symptoms-drug relationship, etc. The challenge with leveraging the authoritative sources to infer answers for non-factoid question lies in: (a) The effective navigation of the answer search-space for timely response to the queries, (b) Ranking the candidate answers derived in step-(a) to enable non-trivial and novel discoveries, and (c) Being robust to perform confirmatory as well as discovery type of tasks.

## 1 Introduction

Hypothesis generation is a crucial element in making scientific discoveries [40, 9, 34, 19, 42, 9, 18, 15, 41, 13, 36, 10, 11, 6, 38, 5, 16, 17, 12, 37, 39, 4, 35]. Traditionally, scientists form hypotheses based on their intuition, ability to make creative chance connections, prior knowledge and experience. This involves them to selectively read hundreds of articles to develop testable hypotheses. However, in the present data-intensive era, it is infeasible for an individual scientist or a research team to keep up with all the relevant articles published in their area of interest. As an example, consider a common disease as *diabetes*, to date (2017) more than 500,000 articles have been published. If a team of scientists peruse around 20 papers per day, it would probably them take 68 years, by which time millions more would have been published [28]. To make things more severe, in domain such as (bio)medicine, hypotheses testing is highly expensive due to the considerable monetary cost associated with clinical experimentation and validation. These problematic issues gives rise to the need for a system that can read, reason, and design hypotheses that transcends the traditional hands-on approach of conducting scientific inquiry. Towards this end, Text Mining, a sub-branch of Data Mining enables researchers to process and analyze natural language text, by sifting through the evidence trails present in multiple dimensions across the corpus and then identifying implicit connections that are revealing but at the same time hitherto unknown. More recently, text mining applied to the domain of biomedicine has been commonly referred to as *conceptual biology* and its importance in fueling hypothesis driven biomedical explorations has been widely recognized [32, 33]. As an example, consider the scenario introduced by Miller et al., [25] - in here the authors are interested in finding a mechanistic link between *Hypogonadism* and Diminished *Sleep Quality* in Aging Men. By applying the ideas from Text mining, in their experiments, the researchers found that *Cortisol* (a steroid hormone) elucidated the observed correlation between hypogonadism and diminished sleep quality in aging men, thereby substantiating a novel hypothesis. Other prominent examples include finding functional connection between genes [2], drug-disease association [33], identification of viruses as bioweapons [32].

Motivated by the potential impact of conceptual biology in the community, we intend to develop a robust computational knowledge discovery framework that enables scientists to make well informed choices on hypotheses testing thereby reducing both the uncertainty involved and the corresponding monetary burden incurred due to invalid experimentation. In this article, we present our initial steps taken towards this direction.

## 2 Problem Statement

Consider two separate literature sets, AL and CL, where the documents in AL discuss concept A and documents in CL discuss concept C. Both of these two literature sets are disjoint. Now, the goal is to find some intermediate concepts B (also known as *bridge* concepts) that connects them in a meaningful way. These bridge concepts are expected to provide novel insights to elucidate the meaning of otherwise poorly understood associations. In the literature, this approach is commonly referred to as the ABC model [32]. Simply put, the ABC model says that if A influences B, and B influences C, then by transitivity A influences C. This derived knowledge or relationship (A implies C) is not conclusive but hypothetical. Further clinical experimentation is required to legitimize it. Over the past decade, practitioners have proposed a range of solutions to find these chains of evidence trails (a.k.a. “hypotheses”). More specifically, as the crux of the problem lies in finding the linking concepts, most of previous solutions tend to focus on the use of distributional statistics, graph theoretic measures and supervised machine learning approaches. However, in broad sense they are afflicted with two major drawbacks. First, these methods assume the prevailing domain to be static, nevertheless it is known that the fields in general (and in particular (bio)medicine) are highly evolving with new facts being added every single day [7]. Thus, for an advanced hypothesis generation tool, it is important to take into account the temporal dynamics of a particular domain. Second, almost all of the previous approaches rely on a pre-defined schema/structure (e.g. graph) that results in finding only those linkages that are en route. As a consequence, it might miss the connections that are surprising or radical. More often, these radical linkages have the potential of shedding novel insights into pathways that remain otherwise hidden. Therefore, a sophisticated hypothesss generation system should be able to capture these intricate complexities of the knowledge diffusion process.

## 3 High-Level Idea

To tackle the aforementioned challenges in generating meaningful hypotheses, we develop a new hypothesss generation framework, namely, “Bridging Evolutionary Medical Concepts (B-Med)”, that is both temporally sensitive and independent of any preordained schema. The tools that we develop for this purpose are entirely general, but to elucidate that it may have immediate practical implications, we are focusing here on the medical domain.

To this effect, the proposed B-Med framework leverages MEDLINE - the most comprehensive literature repository in the field of life sciences and bio-medicine, as the source of information. Next, the key component in hypothesis generation is to find bridge concepts. But, what kind of concepts could be considered as bridges? Intuitively those concepts that have high semantic relatedness to both the start and end concepts (A and C) are promising candidates to be bridges. As an example consider the triplets (Migraine, Pain, Magnesium) and (Migraine, Epilepsy, Magnesium). The first triplet captures a narrower notion of semantic relatedness as the intermediate concept (Pain) is more related to Migraine than Magnesium. In contrast, the second triplet captures a more broader notion of relatedness in a sense that the intermediate term (Epilepsy) is semantically related to both Migraine and Magnesium [31]. In the current problem of interest, the bridge nodes are expected to be semantically related to both the start and end concepts, thus the proposed model focuses on the latter. Upon characterizing the nature of intermediate concepts the next step is to leverage the temporal dynamics of domain. As mentioned before, the medical domain is highly dynamic with new facts being continuously added to its database. This evolving nature causes the semantic meaning of a medical concept to drift over time. Thus, to remain reflect the current state of knowledge, an advanced hypothesis generation system should be able encompass these evolutionary properties of medical concepts. In the next section, we delve into the methodological details of how we incorporate the semantic relatedness and temporal dynamics into our framework.

## 4 Word Embeddings

To capture the notion of semantic similarity and relatedness, we avail a relatively new concepts of Word Embeddings [1] that are a by-product of advances made in the research area of Deep learning. Word Embeddings also known as distributed representations of words or word vectors are unique real-valued vectors for words that are supposed to capture the semantic meaning of a word. They have been experimentally shown to be capable of capturing both the syntactic and semantic properties of words [23, 26]. Word embeddings were initially obtained as a by-product of deep neural language models to represent words. Before the introduction of deep learning into language models, the standard approach to statistical language modeling was based on one-hot representations (one-of-V vectors, i.e., one entry has a value of 1 and the rest are 0s) and N-grams (counting frequencies of occurrences of short text sequences of length up to N). The invention of word embeddings brought two major advantages to representation learning over classic learning algorithms. First, the semantic regularities can be captured in the embedding space and the semantic similarity between words correlates with their distance in the embedding space. Second, word embeddings enable generalization to new combination of text sequences beyond those seen during training (for N-grams that do not use word embeddings, there are *V* ^*N*^ possible combinations, where *V* is the vocabulary size).

Word embeddings are usually learned based on the distributional hypothesis that says “a word is characterized by the company it keeps”. In practice, this can be easily achieved by training a multilayer neural network which predicts the target word in a sequence based on the local context of surrounding words or earlier words. Figure 3 shows an example of the deep language model on a sequence of medical words.Given local context “fish oils can treat”, the target word is constrained to be a kind of disease that can be treated by fish oils, and specifically in this training sequence, the target word is “raynaud disease”. The first layer is an embedding look-up layer where each context word is replaced with a unique word vector. The other layers learn to convert the input word embeddings into the probability distribution for any vocabulary word to appear as the target word. Each dimension of the learned word embeddings can be interpreted as a separate hidden feature of the word, which was not explicitly present in the original text sequences. Since the dimensions (the features) of words are not mutually exclusive and work as a whole to express the complete semantic regularities, word embeddings are also called distributed representations.

Word embeddings are automatically learned by the deep neural language model, rather than manually determined by domain experts ahead of time. Word embeddings are now not only widely used in natural language processing but also are a popular representation learning technique in many research domains, such as bioinformatics and social networks.

In our present system, we learn word vectors from MEDLINE documents that are composed of Medical Subject Headings (MeSH) terms. We call the learned word embeddings for medical terms “MeSH embeddings”. Figure 2 shows an example of the twodimensional projection of MeSH embeddings using t-Distributed Stochastic Neighbor Embedding (t-SNE) [22]. As one can see, MeSH terms with similar medical properties are mapped to the nearby areas in the embedding space.

**Figure 1:**
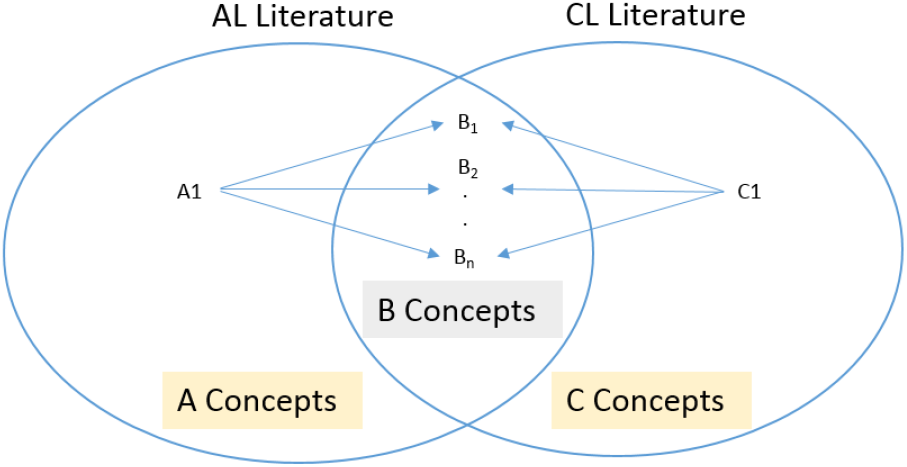
Classical ABC model

**Figure 2:**
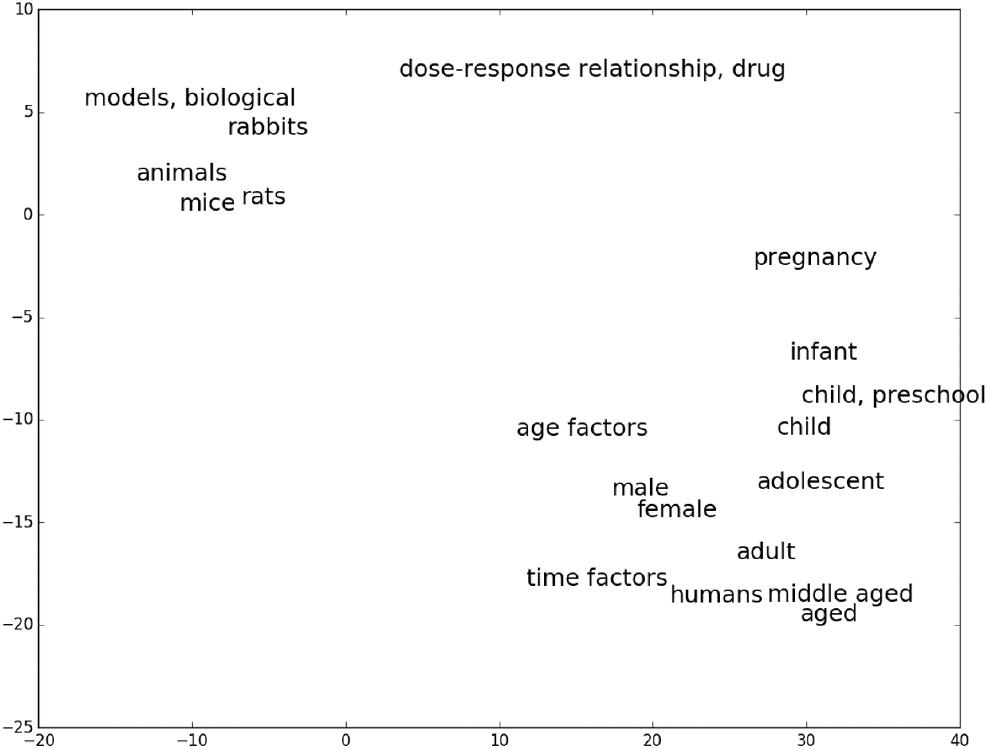
Two dimensional projection of MeSH embeddings

**Figure 3:**
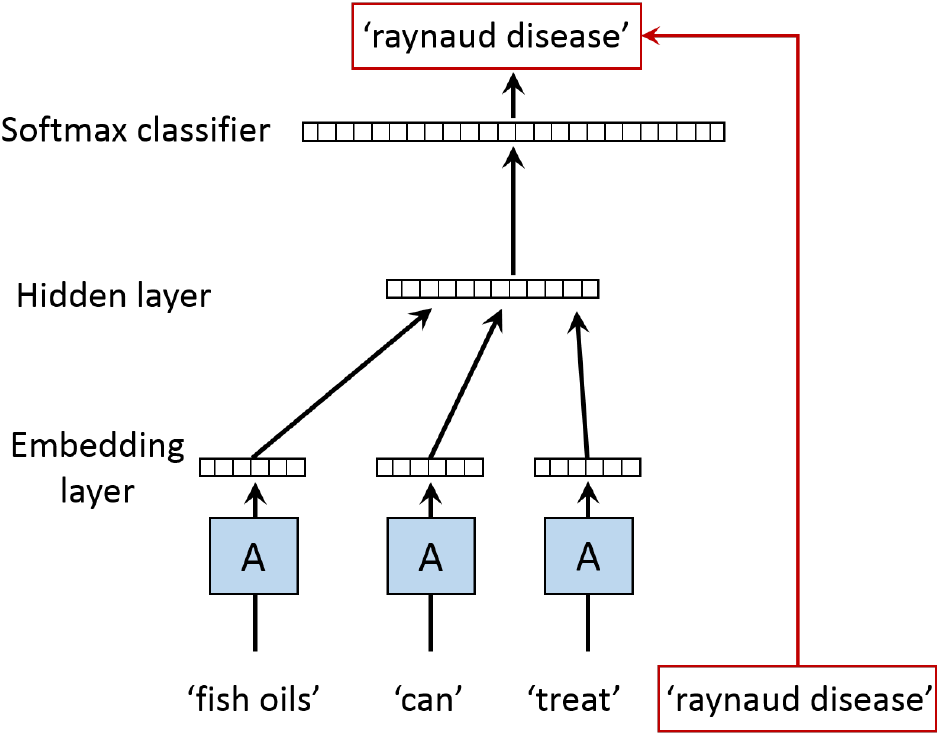
Deep language model

## 5 Proposed Model (Dynamic MeSH Embeddings)

In addition to being able to learn the medical properties of medical concepts (MeSH terms), MeSH embeddings also show promise as a powerful diachronic tool, which is capable of capturing the semantic evolution of MeSH terms [37]. As the body of medical knowledge evolves, for example, the finding of a new treatment or a new cause to a specific disease, the semantics (the medical properties) of the medical concepts evolve too.

In our present B-Med framework, we leverage the MeSH embeddings with an augmented notion of time component to capture the evolutionary properties of medical concepts by using the Dynamic MeSH Embeddings (DME) model. In the dynamic embedding space, the semantic change of a MeSH term can be easily modeled as the location shift of this term. Hence, MeSH terms are projected into the vector space based on their medical properties and gradually drift over time as they evolve. As an example, consider Figure 4, initially during the year of 1985, term “homosexuality” used to be associated with *diseases*, however in recent times (2016) it has evolved to be more related to *gender identity*. This illustrates that the semantic meanings of medical concepts change over the period of time. Thus, to generate semantically sensible hypotheses it is vital to capture the evolutionary characteristics of medical concepts.

**Figure 4:**
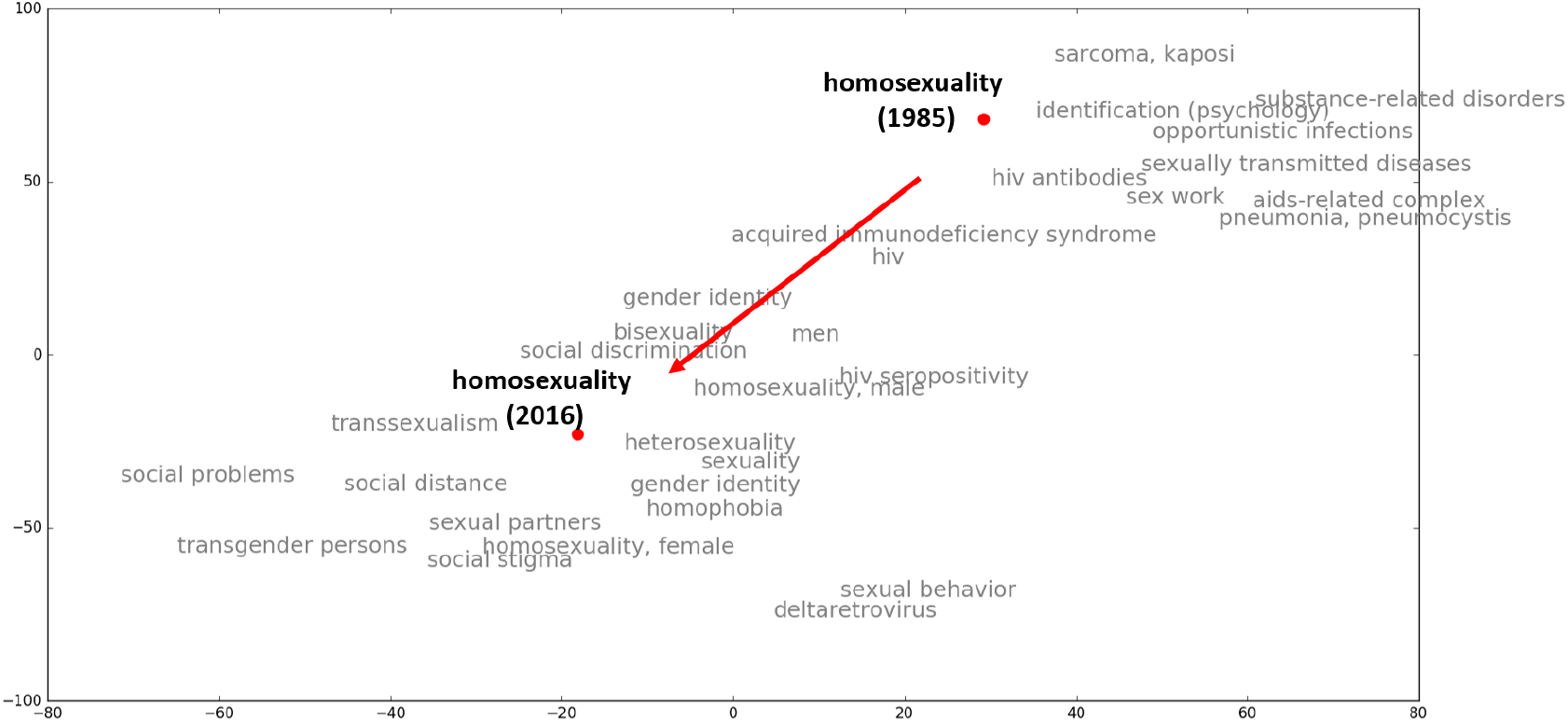
Semantic change of term “homosexuality” over time

Our DME model learns dynamic MeSH embeddings from both the local context in medical documents and the medical concept evolution in history, as shown in Figure 5. The MEDLINE corpus is split into different time slices to provide temporal usage of MeSH terms, for example, 1900-1904, 1905-1909, 1910-1914 and so on.

**Figure 5:**
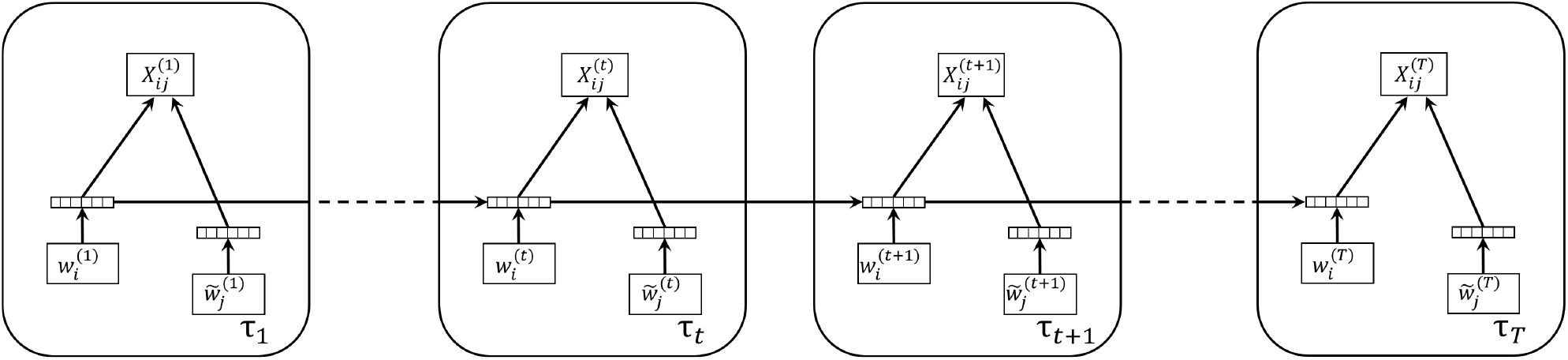
Framework of the DME model

On one hand, our dynamic MeSH embeddings at each time stamp is learned from the local context in medical documents of the corresponding time slice. Specifically, the local context information of MeSH terms is described by the term-term co-occurrence matrix. On the other hand, our dynamic MeSH embeddings learned for each time stamp should also account for semantics present in their historical evolution. To achieve this, at each time stamp, we add a distance constraint to each MeSH term that prevents its embedding from drifting too far from its historical locations.

In this way, temporal component is integrated into MeSH embeddings and smooth evolutionary trajectories of MeSH terms are learned by DME. The association change between any two MeSH terms can be tracked and predicted in the dynamic MeSH embedding space. This provides the possibility to analyze hypothesis generation from temporal aspect.

## 6 Hypothesis Generation

The learned dynamic MeSH embeddings enable us to generate medical hypotheses that are both temporally sensitive and semantically meaningful. Given two MeSH terms *i* and *j* that do not have any direct associations before time *t*, a hypothesis (bridge term) *k* refers to another MeSH term which has a high probability to connect *i* and *j* in a future article. For example, term *i* is a disease, term *j* is a kind of drug, and bridge term *k* might denote the mechanism how *j* treats *i*. Intuitively, a good hypothesis should be semantically related to both the inquiry terms and has an evolutionary trend towards the direction that the inquiry terms evolve. Moreover, general terms such as “human” that is semantically related to several other terms terms, are not good hypothesis terms as they are not informative.

More specifically, when generating hypotheses *k* for inquiry terms *i* and *j* at time *t*, three factors are taken into consideration. First, we examine term *k*’s cosine similarity with terms *i* and *j* in the vector space at time *t*, to evaluate the current semantic association strength between them. As demonstrated in many previous studies, the cosine similarity between two word embeddings correlates with their semantic similarity [1, 24, 21]. This ensures that the generated hypothesis has coherent medical properties with the input inquiry terms. Second, we analyze the evolution trends of the terms by tracking their evolutionary trajectories in the dynamic embedding space. Hypothesis term *k* is favored if there is a growing trend of association strength between *k* and inquiry terms *i* and *j*, in other words, an increasing trend of average cosine similarity between *i, j* and *k* over time. With an increasing evolutionary trend towards the direction the inquiry terms evolve, the generated hypothesis has a higher chance to form a connection with them in the future. Third, we promote informative bridge terms and demote general terms when ranking hypotheses by checking their evolutionary behavior over time. According to the law of conformity reported in [8] – “rates of semantic change scale with a negative power of word frequency”. This is consistent with our intuition and observations. General terms with high term frequency, such as “human”, “male” and “female”, have very stable semantics, manifested as being geographically stable in the dynamic vector space over time. In contrast, informative bridge terms, such as “homosexuality”, may shift locations in the dynamic vector space over time as their semantics change and our biomedical knowledge evolve. Hence, by analyzing the evolutionary dynamic MeSH embeddings, we are able to generate hypotheses that are informative, temporally sensible and semantically coherent with the inquiry terms.

We demonstrate the efficacy of our model on a set of real-world hypothesis generation test cases. It should be noted that evaluating hypothesis generation systems is not a simple task and remains an open problem [43]. This is due to the unavailability of a comprehensive ground truth dataset of validated hypotheses, and the lack of a systematic methodology to compare different rankings of generated hypotheses. To overcome this difficulty, replicating existing scientific discoveries based on historical biomedical literature has been seen as an effective approach to evaluate generated hypotheses by most researchers. The pioneers in this area of study Swanson and Smalheiser [30] proposed their initial hypothesis generation model and published several biomedical hypotheses based on the literature, which were subsequently clinically validated. Since then, these proposed terms in their discoveries have become a gold standard for evaluating hypothesis generation models. The followings are the *de facto* gold test cases that have certain expected results:

1. Fish-oil (FO) and Raynaud’s Disease (RD) (1985)
2. Magnesium (MG) and Migraine Disorder (MIG) (1988)
3. Somatomedin C (IGF1) and Arginine (ARG) (1994)
4. Indomethacin (INN) and Alzheimer Disease (AD) (1989)
5. Schizophrenia (SZ) and Calcium Independent Phospholipase A2 (CI-PA2) (1997)

As one can see in each test case, apart from two input terms, there is a cutoff date indicating the first time the two inquiry terms co-occur in literature. Therefore, to assess the effectiveness of our model, for each test case, we will only use the dynamic MeSH embeddings before the cutoff date to generate hypothesis.

Figure 6 shows three examples of how our dynamic MeSH embeddings facilitate the process of hypothesis generation. Figure 6.(a) shows the two-dimensional projection of the evolutionary trajectories of FO and RD in gold test case 1 and their bridge term Blood Viscosity (BV) before the cutoff date using t-SNE. There was no direct associations between FO and RD initially until they were connected by BV in 1985. As one can observe, initially in 1953, all three MeSH terms were far apart from each other, but as biomedical researchers continuously studied over these topics in parallel, their implicit semantics (medical properties) started to get closer and closer to each other. This evolutionary behavior eventually in the year of 1986 led to their very first co-occurrence in a research article. This growing association trend among these MeSH terms can be quantitatively measured by the average cosine similarity in the dynamic embedding space. Figure 6.(b) and Figure 6.(c) illustrate the evolutionary behavior of gold test cases 2 and 4, respectively. As with test case 1, the inquiry terms and the ground truth bridge terms also show an increasing trend of semantic similarities before the association between them is formed.

**Figure 6:**
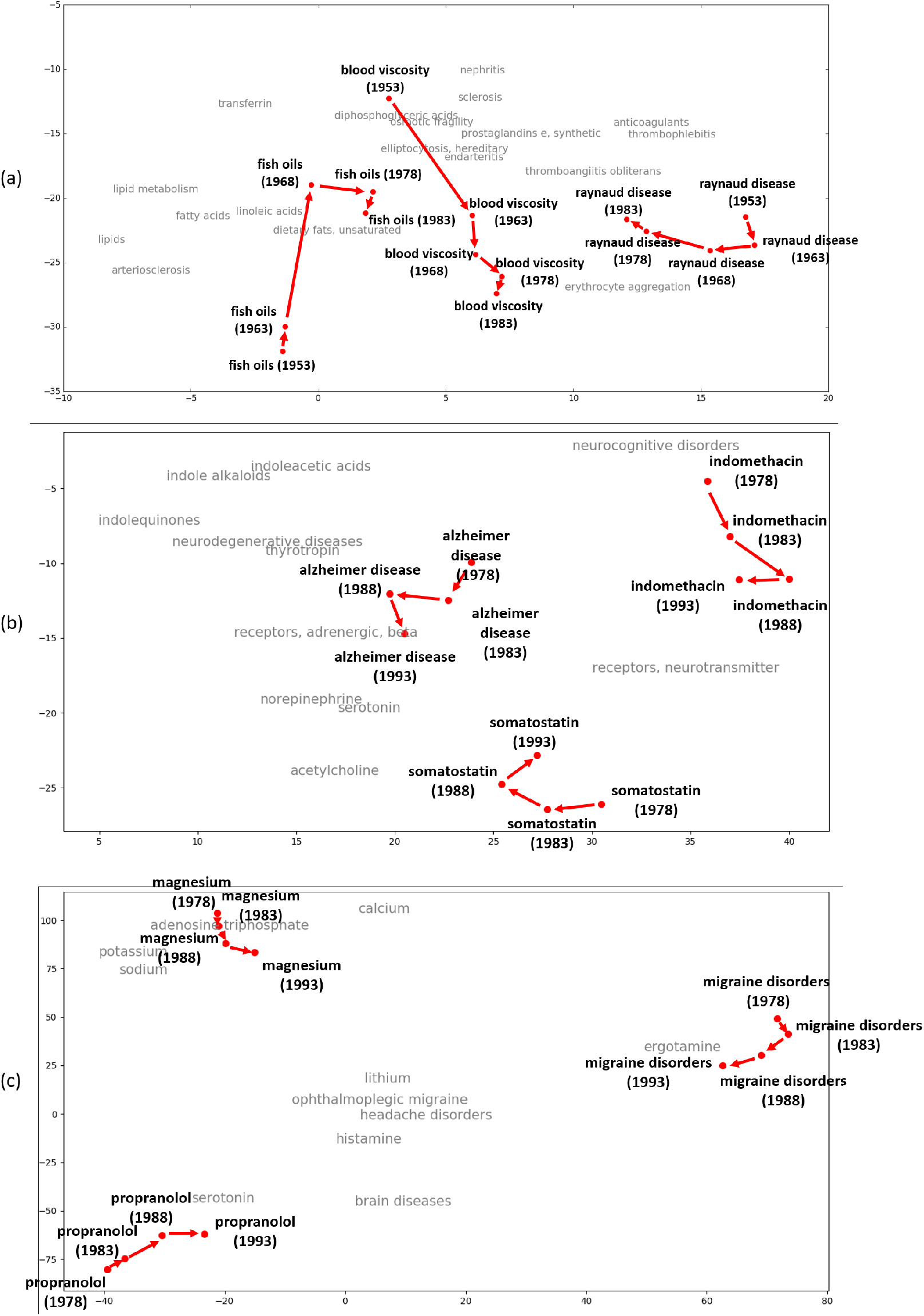
Evolutionary trajectories for the gold test cases

In Table 1, top 15 generated hypotheses for each of the gold test cases are reported. We are interested to see if our model is capable of rediscovering already established biomedical knowledge based on historical knowledge, as well as to examine the validity of other high ranked bridge terms. The evaluation is done by comparing our ranked hypothesis list in Table 1 with the *de facto* gold results and manually checking the relationship between the generated hypotheses and the inquiry terms in the biomedical literature. To get the medical journal for manual inspection, we formulate a boolean query on Google and examine top 15 results, for example, fish oils AND epoprostenol AND raynaud’s disease. For each of the valid hypothesis in Table 1, we provide its corresponding PubMed identifier (PMID). A hypothesis is considered to be a valid connection if it co-occurs together with the input terms after the cut-off date.

**Table 1.**
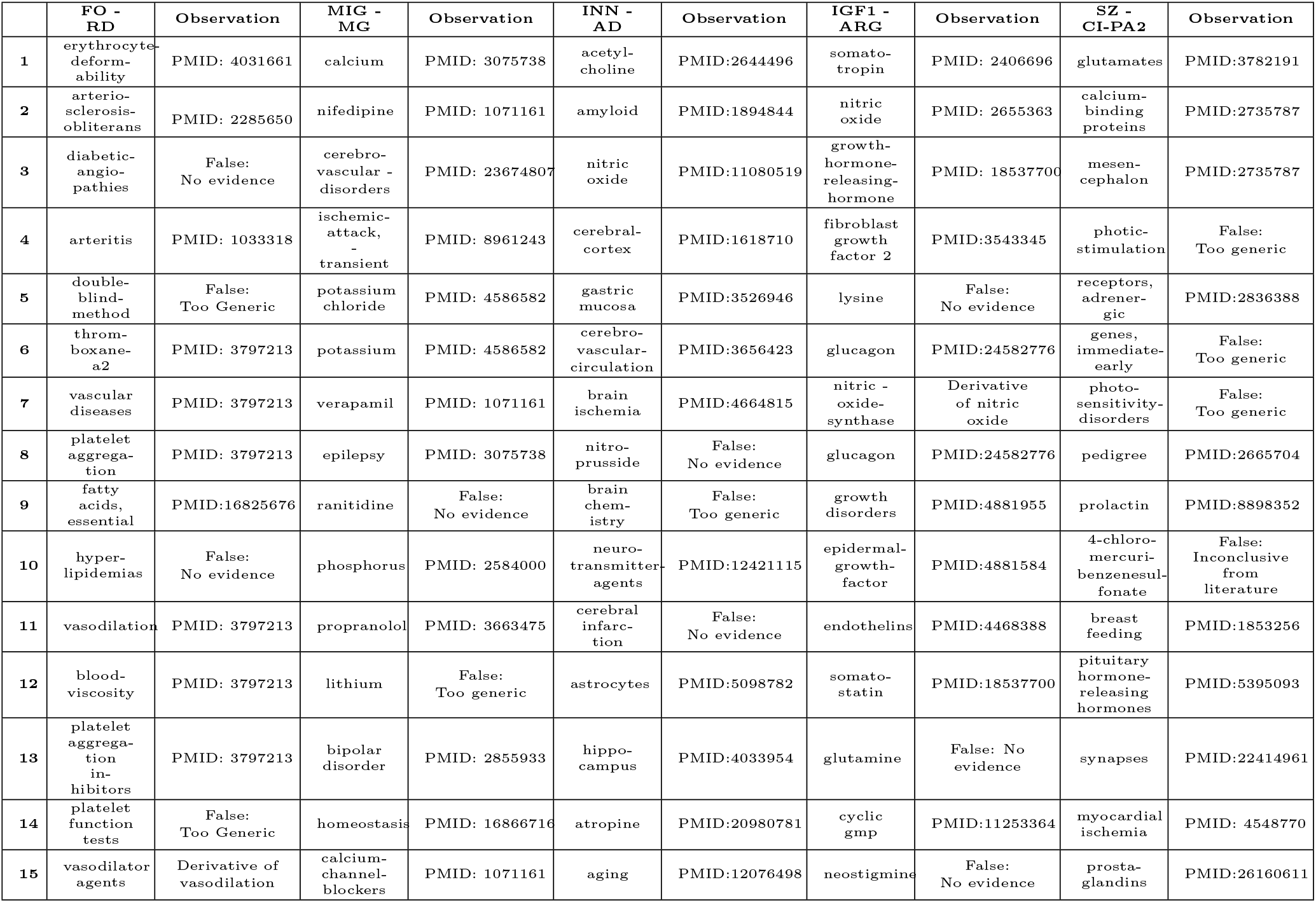
Top 15 Intermediary MeSH Terms for Each Test

|  | FO - RD | Observation | MIG - MG | Observation | INN - AD | Observation | IGF1 - ARG | Observation | SZ - CI-PA2 | Observation |
| --- | --- | --- | --- | --- | --- | --- | --- | --- | --- | --- |
| 1 | erythrocyte-deformability | PMID: 4031661 | calcium | PMID: 3075738 | acetylcholine | PMID:2644496 | somatotropin | PMID: 2406696 | glutamates | PMID:3782191 |
| 2 | arteriosclerosis-obliterans | PMID: 2285650 | nifedipine | PMID: 1071161 | amyloid | PMID:1894844 | nitric oxide | PMID: 2655363 | calcium-binding proteins | PMID:2735787 |
| 3 | diabetic-angiopathies | False: No evidence | cerebro-vascular - disorders | PMID: 23674807 | nitric oxide | PMID:11080519 | growth-hormone-releasing-hormone | PMID: 18537700 | mesencephalon | PMID:2735787 |
| 4 | arteritis | PMID: 1033318 | ischemic-attack, -transient | PMID: 8961243 | cerebral-cortex | PMID:1618710 | fibroblast growth factor 2 | PMID:3543345 | photic-stimulation | False: Too generic |
| 5 | double-blind-method | False: Too Generic | potassium chloride | PMID: 4586582 | gastric mucosa | PMID:3526946 | lysine | False: No evidence | receptors, adrenergic | PMID:2836388 |
| 6 | thromboxane-a2 | PMID: 3797213 | potassium | PMID: 4586582 | cerebro-vascular-circulation | PMID:3656423 | glucagon | PMID:24582776 | genes, immediate-early | False: Too generic |
| 7 | vascular diseases | PMID: 3797213 | verapamil | PMID: 1071161 | brain ischemia | PMID:4664815 | nitric - oxide-synthase | Derivative of nitric oxide | photosensitivity-disorders | False: Too generic |
| 8 | platelet aggregation | PMID: 3797213 | epilepsy | PMID: 3075738 | nitro-prusside | False: No evidence | glucagon | PMID:24582776 | pedigree | PMID:2665704 |
| 9 | fatty acids, essential | PMID:16825676 | ranitidine | False: No evidence | brain chemistry | False: Too generic | growth disorders | PMID:4881955 | prolactin | PMID:8898352 |
| 10 | hyper-lipidemias | False: No evidence | phosphorus | PMID: 2584000 | neuro-transmitter-agents | PMID:12421115 | epidermal-growth-factor | PMID:4881584 | 4-chloro-mercuri-benzenesulfonate | False: Inconclusive from literature |
| 11 | vasodilation | PMID: 3797213 | propranolol | PMID: 3663475 | cerebral infarction | False: No evidence | endothelins | PMID:4468388 | breast feeding | PMID:1853256 |
| 12 | blood-viscosity | PMID: 3797213 | lithium | False: Too generic | astrocytes | PMID:5098782 | somato-statin | PMID:18537700 | pituitary hormone-releasing hormones | PMID:5395093 |
| 13 | platelet aggregation inhibitors | PMID: 3797213 | bipolar disorder | PMID: 2855933 | hippocampus | PMID:4033954 | glutamine | False: No evidence | synapses | PMID:22414961 |
| 14 | platelet function tests | False: Too Generic | homeostasis | PMID: 16866716 | atropine | PMID:20980781 | cyclic gmp | PMID:11253364 | myocardial ischemia | PMID: 4548770 |
| 15 | vasodilator agents | Derivative of vasodilation | calcium-channel-blockers | PMID: 1071161 | aging | PMID:12076498 | neostigmine | False: No evidence | prostaglandins | PMID:26160611 |

### 6.1 Fish-oil (FO) and Raynaud’s Disease (RD)

In 1986, Swanson [30] explored the research question of “role of dietary fish oils in treating patients with Raynaud’s syndrome”. Upon manual inspection of literature related to Fish oils and Raynaud’s disease respectively, he found that Raynaud’s disease is aggravated by high *blood viscosity*, high *platelet aggregation, Vasoconstriction*, and the ingestion of Fish oils can reduce these phenomena. As one can observe, in Table 1 both *platelet aggregation* and *blood viscosity* are found at rank 8 and 11 respectively. In addition to these important connections, other terms in our ranked set such as *‘fattyacids, essential’, ‘vasodilation’* are also meaningful pathways.

### 6.2 Magnesium (MG) and Migraine Disorder(MIG)

Swanson [31] proposed eleven hypothetical pathways (bridge terms) for Magnesium and Migraine Disorder. These connections are *epilepsy, serotonin, prostaglandins, platelet aggregation, calcium antagonist, type A personality, vascular tone and reactivity, calcium channel blockers, spreading cortical depression, inflammation, brain hypoxia and substance P*. As can be observed in Table 1, important hypotheses are successfully generated, such as *epilepsy, calcium channel blockers, adenosine triphosphate*, etc. In this regard, it should be noted that previous research indicates this to be a difficult test case [29].

### 6.3 Somatomedin C (SMC) and Arginine (ARG)

Somatomedin C (SMC), also known as Insulinlike Growth Factor I (IFG1), is a growth regulating peptide. Arginine is an important amino acid. Growth hormones such as *somatotropin* and *somatostatin* are found to be valid bridge terms to connect them. ARG stimulates the secretion of growth hormones and growth hormones have an influence in SMC. As it can be seen, somatotropin is ranked as the highest hypothesis by our model and somatostatin is generated in top 15.

### 6.4 Indomethacin (INN) and Alzheimer Disease (AD)

Alzheimer Disease (AD) is a progressive disease that destroys memory and other important mental functions. The research question of whether AD can be treated with an inflammatory agent - Indomethacin (INN) was explored during the 1990s. Researchers reported that Acetylcholine, Membrane fluidity are important bridge terms. In our results, as with the SMC-ARG test case, Acetylcholine is labeled are the most proable hypothesis by our model. In addition, the derivatives of Membrane fluidity were also found in top 15. An interesting observation that we would like to discuss here is regarding term *nitric oxide* (Rank=3). Although not yet validated, several papers identified nitric oxide as important for understanding Alzheimer’s disease [29]. Moreover, during 2000-2001, there were studies [3] showing evidence of strong influence of nitric oxide in both Alzheimer’s disease and Indomethacin.

### 6.5 Schizophrenia (SZ) and Calcium - Independent Phospholipase A2 (CI-PA2)

Schizophrenia is a disorder that affects a person’s ability to think, feel, and behave clearly. It is found that CI-PA2 is elevated in SZ patients. After mannualy examining independent works of [27] and [20], Swanson and Smalheiser postulated oxidative stress to be the key bridge term between SZ and CI-PA2. In our results, we were able to rediscover oxidative stress indirectly through receptors, adrenergic (PMID: 3820966). Also, similar to the previous test case, our topmost ranked term glutamates has been heavily investigated for its influences in treating Schizophrenia (PMID: 20686195) in more recent years.

## 7 Novel Applications

Because of its ability to capture temporal dynamics, the methodological innovation of DME has a broad range of utility in both content analysis and semantic search based applications. To illuminate its swift benefits, in this section, we describe its application in the task of Ontology extension (a sub-problem of Ontology evolution). Generally speaking, Ontologies refer to a formal, axiom based structures such as thesauri, taxonomies, is-a-hierarchies, that aid in consistent exchange of information between entities. Traditionally, the ontologies are maintained by subject matter experts. Being manually curated, these are highly precise, however, the process of maintaining them is often slow, tedious and prone to cumbersome issues. Thus, recent times have witnessed a significant attention from researchers to automate this process. This problem becomes of utmost importance in domain such as bio-medicine where new discoveries are published everyday, bringing into light new facts and rending some obsolete. Moreover, biomedicine is known to harbor abundant ontologies thus requiring them to be regularly updated so as to reflect the new knowledge. This dynamic nature of task makes the concept DME of a suitable choice. To illustrate the problem further, consider Figure 7 as an example, the snapshot shows the evolution of medical concepts^1^ over time in ontology. Figure 7 (a) depicts the snapshot of a medical taxonomy in year 1999. As it can be observed in the succeeding year (2000) (Figure 7(b)) the medical concepts “Abdomen” and “Anostraca” expanded whereas the heading “Mesentery” remained intact. This is the core element in the task of Ontology extension. Given the past ontology versions and related dynamic MeSH embeddings, can we predict the branches in the hierarchy that will undergo expansion in the future (See Figure 7(c)) ?

**Figure 7:**
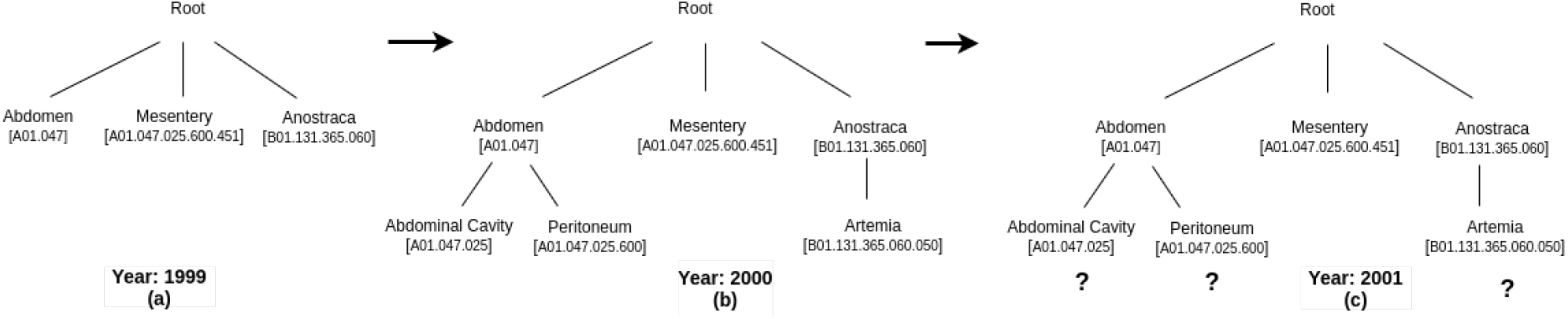
Snapshot depicting evolution of MeSH hierarchy

To tackle this problem, we leverage the methodological benefits of DME. Since the focus of DME is in incorporating the temporal features, we first bring to light the crucial elementary factor that causes a medical concept to expand over time - semantic density change of a medical concept. The semantic density of a medical concept refers to the total aggregation of semantics associated with a concept, manifested as the compactness in its neighborhood region in the embedding space. As the body of medical knowledge evolves the semantic meaning of a medical concept starts to get associated with more and more concepts. Simply put, the number of near-synonyms and subsets of a term in their neighbourhood region increases, causing the area to become denser and denser. For illustration, consider Figure 8, which shows the semantic density change of a medical concept “p38 Mitogen-Activated Protein Kinases”. The red dot represents this medical concept and the black dots represent its top 10 closest neighbor terms in the embedding space. As it can be observed, over the passage of time, the region around the concept becomes denser and denser. We assume this escalation in semantic density gives rise to the need for a higher level of specificity, thus resulting in concept expansion in knowledge ontology.

**Figure 8:**
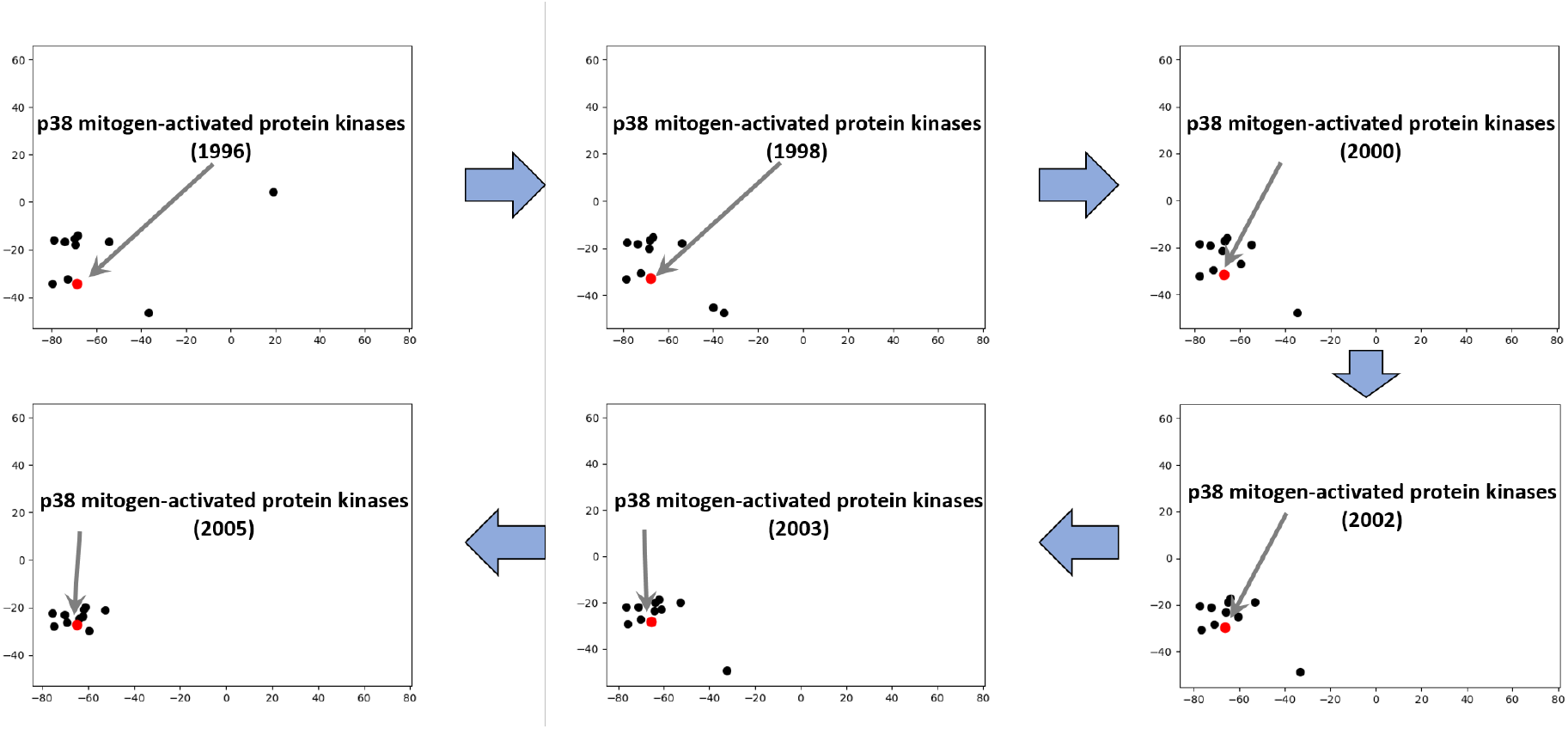
Evolving semantic density of a medical concept.

As DME are adept at incorporating the temporal dynamics, they are our natural choice to measure the semantic density change of MeSH terms. To recapitulate, DME’s are time-aware learned embeddings that at any time *t* are sensitive to the medical semantics of both the current time slice and their diachronic period. As a results, they can capture both the evolving properties of medical concepts and track their semantic progression, thereby, successfully aiding to predict the concepts that will undergo expansion in future. Further methodological details are not within the scope of present article. Interested readers may refer to the article [14]. In the results we found that the proposed methodology is able to predict concept expansion with high accuracy. By precisely pinpointing the areas of ontology that are more likely to undergo extension in the near future, we intend to ease the burden of manually keeping an ontology up-to-date for the domain experts, thus allowing them to focus on more complex ontology evolution issues.

## 8 Summary and Future Directions

An advanced hypothesis generation system that generates “actionable” postulates is a particularly difficult task, given the intrinsic complexities present in the process of imitating the steps a cognitive mind undertakes while forging a plausible hypothesis. However, the massive amount of data being generated by health-care sector and its current trend towards rapid digitization has overwhelmed the domain experts. Consequently, it has become necessary to design a system that can process, analyze this quintessential (bio)-medical “big data” and generate promising hypotheses that could be potentially validated, thereby, benefiting the society at large.

In this direction, the proposed B-Med framework is our initial step towards developing a robust hypothesis generation system. Aside from the evolutionary characteristics it carries and the flexibility it provides in integrating heterogeneous textual sources, it is computationally tractable. This allows the end users to obtain the desired results in a reasonable amount time. Furthermore, another crucial advantage of this framework lies in its ability to present evolution trajectory visualization of medical concepts. Similar to Figure 6, for any two medical concepts, their semantic progression over time can be analyzed and understood. This form of visualization is believed to aid domain experts in making informed choices. Finally, the system also provides evidences (PubMED article identifier) for its outputs thus making its results interpretable.

The results indicate that the hypotheses generated by the proposed B-Med framework are encouraging. As an illustration, consider the bridge concept *nitric oxide* that was found for the test-case of Alzheimer’s disease and Indomethacin. Although not clinically corroborated yet, several papers identified nitric oxide as an important element for understanding Alzheimer’s Disease [29]. Additionally, during 2000-2001, there were studies [3] showing evidence of strong influence of nitric oxide in both Alzheimer’s disease and Indomethacin. Thus, the results suggest that the proposed methodology is capable of generating both semantically meaningful and temporally sensible hypotheses that are worthy of clinical trails and further investigation.

In our continuing research, we are investigating in several directions. First is to speculate more sophisticated approaches to generate MeSH embeddings that are able to capture the multifaceted aspects of semantic expressiveness. Another area of interest is to explore the application of our methodology to tasks such as drug-drug interaction, adverse-drug events and biomedical question answering.

## Footnotes

1 https://www.nlm.nih.gov/pubs/factsheets/mesh.html

## Notes

### Competing Interest Statement

The authors have declared no competing interest.

